# Bioassay evaluation, GC-MS Analysis and Acute Toxicity Assessment of Methanolic Extracts of *Trichoderma* species in an Animal Model

**DOI:** 10.1101/2022.02.15.480498

**Authors:** Afrasa Mulatu, Negussie Megersa, Teshome Tolcha, Tesfaye Alemu, Ramesh R. Vetukuri

## Abstract

Fungi of the genus *Trichoderma* have been marketed for the control of crop pests, weeds, and diseases. However, some *Trichoderma* species may produce toxic secondary metabolites; the existence of these compounds in bioformulated products, along with their relative risk, should receive due attention to ensure human safety. In this study, we investigated the in vitro antagonistic potential of *T. asperellum* AU131 and *T. longibrachiatum* AU158 as a biocontrol agent against *F. xylarioides* and the associated antagonistic mechanism with bioactive substances. Swiss albino mice were used to evaluate the in vivo toxicity and pathogenicity of *T. asperellum* AU131 and T. *longibrachiatum* AU158 methanolic crude extracts and spore suspensions, respectively, in a preliminary safety assessment for use as biofungicides. Metabolite profiling of the crude extracts were performed by Gas Chromatography-Mass Spectrometry (GC-MS). The agar diffusion assay of the crude extracts from both *T. asperellum* AU131 and *T. longibrachiatum* AU158 were the most effective at a concentration of 200μg/mL, causing 62.5%, and 74.3% inhibition, respectively. A GC-MS analysis of crude extracts from both bioagents identified 23 volatile secondary metabolites classified as alcohols, acids, sesquiterpenes, ketones and aromatic compounds. The oral administration of crude extracts (doses of 600, 1200, 2000 and 5000 mg/kg body weight) and spore suspensions (doses of 1 × 10^6^, 1 × 10^7,^ 1 × 10^8^ and 1 × 10^9^ spores/mL) to female Swiss albino mice over 14 days did not show any significant signs of toxicity, mortality or changes to body weight. The highest dose, 5000 mg/kg, can be used to determine the no observed effect level of crude extracts and spore suspensions of *T. asperellum* AU131 and *T. longibrachiatum* AU158 when calculating safety margins. It can be concluded that these two *Trichoderma* species can serve as a biocontrol agents against *F. xylarioides* and the mechanism for this function was due to the secondary metabolites with effective bioactive substance. Moreover, the tested spore suspensions and crude extracts were not pathogenic or toxic, respectively, when administered to Swiss albino mice at various doses.

## 1. Introduction

Given the large socio-economic impact of crop monocultures and the environmental hazards of chemical pesticides [1], biocontrol agents (BCAs) have recently become a strategic option for controlling plant diseases and enhancing crop yields [2]. A number of biopesticides that contain fungal species as the active ingredients are currently marketed in Europe, USA, Asia and Africa [3]. In contrast to chemical pesticides, which have been reported to induce resistance among pests and cause residual toxic effects, biopesticides are highly selective against the target pest [4]. Biocontrol agents have been designed to grow and reproduce, surviving in the environment for prolonged periods in symbiotic consortia with the host; this means that relative to chemical pesticides, only a small amount of bioagents needs to be applied to a certain location [5,6]. Although the vast majority of BCAs are generally regarded as safe for humans and the environment [7], some studies have demonstrated that increased exposure to fungal substances among agricultural workers may affect the immune system [8].

There is evidence that members of the genus *Trichoderma* are effective biocontrol agents against various plant pathogens across different agro-ecological systems [6,9-12]. However, *Trichoderma* species are also known to cause opportunistic infections in humans, varying from localized to fatal disseminated diseases; these are particularly dangerous for risk populations, including patients undergoing peritoneal dialysis, transplant recipients and patients with hematological malignancies [13,14]. In the previous study, we reported a comparatively high *Trichoderma* species diversity in coffee ecosystem in Ethiopia among which *T. asperellum* AU131 and *T. longibrachiatum* AU158 were found effective against *F. xylarioides* [15]. Thus, we formulated a biofungicide from *T. asperellum* AU131 and *T. longibrachiatum* AU158 for the control of coffee wilt disease (CWD) caused by *F. xylarioides* [16]. Subsequent studies demonstrated that both species benefit the coffee plant by inhibiting the development of *F. xylarioides* in the rhizosphere. For this reason, systematic studies are necessary for evaluating and ensuring the safety of biofungicides for agricultural use [17]. On the other hand, understanding and evaluating the mechanism(s) of secondary metabolites of these potential biocontrol agents against *F. xylarioides* is very important. Several *Trichoderma* strains were widely studied due to their capacity to compete for nutrients and space [18], parasitize other fungi [19,20], enact antibiosis by producing secondary metabolites or antimicrobial compounds [8,21,22], induce defense responses in plants [17,18], and promote plant growth [15,19]. *Trichoderma* species produce and emit secondary metabolites associated with antimicrobial ability, induce defense responses in plants, and promote plant growth [23].

Since the 1960s, toxicity studies have been used as a vital step in the approval of new products onto the market. The present study was conducted in animal models following the protocols laid out by the Organization for Economic Cooperation and Development (OECD) [24]. More specifically, the study applied the experimental design described in Tier 1 of the Toxicological Evaluation of Microbial Pest Control Agents (MPCA), which has the purpose of assessing the pathogenicity and toxicity of products [25]. In Europe, Council Directive 91/414/EEC and its successive amendments identify which requirements an applicant must achieve for the authorization to produce and market pesticides, including those that have a BCA as an active substance [26,27]. In particular, the directive requires the provision of information concerning the short-term toxicity of any relevant metabolites produced by the candidate BCA [27-29]. The present study provides toxicological data that are of importance in assessing the toxicological relevance of two potential BCAs produced by *T. asperellum* AU131 and *T. longibrachiatum* AU158. Therefore, it is conceivable that it would be more effective to determine the toxicological risks associated with a particular BCA by assaying a mixture of metabolites, like those in crude extracts, on model animals that are sensitive to a large spectrum of molecules, instead of assessing the toxicity of single and pure metabolites [26,27].

In the present study, Swiss albino mice were used to assess the acute toxicity of spore suspensions and bioactive methanolic extracts of *T. asperellum* AU131 and *T. longibrachiatum* AU158 in short term assays. Thus, the study aimed: (i) to evaluate the sensitivity of albino mice to secondary metabolites produced by both *Trichoderma* species in preliminary toxicological screenings; (ii) to generate toxicological data that can be used when assessing the risk associated with toxicity of metabolites produced by both *Trichoderma* species; (iii) to evaluate the in vitro antagonistic effect of these secondary metabolites against *F. xylarioides* and (iv) to profile volatile secondary metabolites produced by these fungi through Gas Chromatography-Mass Spectrometry (GC-MS) analysis.

## 2. Materials and Methods

### 2.1 Trichoderma species and preparation of spore suspensions

In our previous work, *T. asperellum* AU131 and *T. longibrachiatum* AU158 were used in the development and formulation of biofungicides for the control of CWD, which is caused by *Fusarium xylarioides* [16]. When studying acute oral toxicity, spore suspensions (live cells) were prepared by adding 5 mL (0.9%) of NaCl to mature *Trichoderma* species on PDA plates to dislodge the spores from the mycelium [30]. The spores were counted using a hemocytometer (Neuberger GmbH, Germany) to determine the spore concentrations.

### 2.2 Effect of temperature and pH on mycelial growth

To evaluate the virulence of the biocontrol agents, mycelial growth at different temperatures, from 25 to 37°C, were inoculated onto Petri dishes containing minimal (0.5% glucose, 0.1% KH_2_PO_4_, 0.1% (NH_4_)_2_SO_4_, 0.1% MgSO_4_.7H_2_O, 2% agar in distilled water) or yeast extract (0.5% glucose, 0.2% yeast extract, 1% KH_2_PO_4_, 2% agar in distilled water) agar medium. The effect of pH on mycelial growth was determined by using buffer solutions to adjust the pH of both minimal and yeast extract agar medium to values ranging from 2 to 9 [13]. Mycelial discs (5 mm in diameter) cut from the active growing culture were used as inocula. After incubation at different temperatures, mycelial growth was determined by colony diameter measurements.

### 2.3 Extraction of secondary metabolites

For secondary metabolite extraction, *Trichoderma* species were grown in 500 mL flasks containing 100 g (dry weight) solid substrate (wheat bran to white rice (2:1 *w/w*)) supplemented with 1% (v/v) glycerol and 1% (w/v) (NH_4_)_2_SO_4_) to moisten the substrate [16]. The flasks were sterilized at 121°C for 15 min and, after cooling, inoculated with 20 mL spore suspension (5 × 10^7^ spores/mL) followed by incubation at 28°C for 21 days. The cultures were homogenized by adding methanol, followed by centrifugation at 5000 rpm for 15 min. The supernatant was collected and dried under a vacuum; the obtained crude extract was purified using Sephadex LH-20 column (Sigma-Aldrich, St. Louis, MO) and stored at -20°C until use for oral administration in mice and GC-MS analysis.

### 2.4 Toxicological bioassay of secondary metabolites against F. xylarioides

#### 2.4.1 Seeded agar assay

The methanolic crude extract of *T. asperellum* AU131 and *T. longibrachiatum* AU158 were filtered through 0.22 μm nylon membranes to get purified extracts. The crude extracts were mixed with warm SFM agar (200, 100, 50 and 10μg/mL of crude extracts) and plated in sterile Petri plate to analyze the effect of non-volatile secondary metabolites on the test pathogen [31]. A 5 mm mycelial disc of the test pathogen was placed at the center of the Petri plate-containing SFM agar. Plates without crude extract was served as control. These plates were maintained at 28°C until the control plate was fully-grown. The diameter (mm) of the fungal colony growth was measured, and the percent inhibition (PGI) was calculated using the formula,

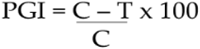

Where, PI = percent inhibition; C = radial growth of the pathogen in control plate; T = radial growth of the pathogen in treatment.

#### 2.4.2 Agar diffusion assays

To evaluate antifungal activity of the crude extract, 1 mL of spore suspension (5 × 10^5^ spore/mL) of *F. xylarioides*, was spread on the surface of the plate containing King B (KB) medium and 5 mm diameter wells were bored into the agar circularly [32,33]. Fifty μL of the series dilution of the crude extract was poured into the well and methanol was used as negative control. The plates were incubated at 25°C for 3 days. Three parallel experiments were set up to measure the inhibition zones. The inhibition zones were measured as the diameter of the fungal and bacterial colonies, and was expressed as the percentage of growth inhibition (PGI). Inhibition zone = average diameter of the colony-5 mm (diameter).

### 2.5 GC-MS metabolite profiling of Trichoderma species crude extracts

The volatile metabolites were analysed by gas chromatography with a single quadrupole mass spectrometer detector (GC-MS) analysis [34,35]. Prior to the analysis, dried crude extracts obtained from *Trichoderma* species were dissolved in ethyl acetate and placed in a 1 mL glass vial. The GC system was equipped with an HP-5MS (30 m × 0.25 mm and 0.25 μm 5% diphenyl/95% dimethylpolysiloxane) capillary column (J and W Scientific, Folsom, CA, USA). The instrument was programmed to start at 40°C (held for 2 min) and to increase to 160°C by 6°C /min, after which it increased to 260°C by 10°C/min (held for 4 min). Helium (99.9%) was used as the carrier gas, and the flow rate was 1 mL/min. The flow was then transferred from the column to an Agilent 5973 MS detector (Agilent Technologies, Palo Alto, CA). The ion source temperature was set at 230°C, the ionizing electron energy was 70 eV, and the mass range was 40–450 Da in full-scan acquisition mode [33]. The spectra of identified metabolites were compared with the spectra of known compounds in the GC-MS NIST database. The threshold for identification was ≥90% similarity. The analysis was carried out in triplicate to monitor the repeatability of the analysis. The instrumental responses obtained were interpreted using mass hunter ChemStation software.

### 2.6 Experimental animal and Ethical approval

Adult female Swiss albino mice (6-8 weeks old), nulliparous, non-pregnant and weighing 25-30 g [29,36], were obtained from the Ethiopian Public Health Institute (Addis Ababa, Ethiopia). Female mice were allocated in treatment groups (five animals/per group). All of the animals were kept in polypropylene cages and acclimatized for a period of seven days prior to the start of the experiment. They were placed under standard conditions (23 ± 2°C with a 12 h light-dark cycle), and were fed commercial rodent chow *ad libitum* and given clean tap water during the experimental period [17]. All of the animals were cared for in compliance with the internationally accepted guide for the care and use of laboratory animals [37], as well as Addis Ababa University institutional guidelines for animal ethics, which were previously approved by the College of Natural and Computational Sciences Institutional Review Board (CNCS-IRB) under the protocol number IRB/44/2020.

### 2.7 Experimental Design

For the effective administration of crude methanolic extracts and spore suspensions to mice through oral gavage, water was not provided in the morning to induce thirst. After fasting for 3 h, crude extracts and live cells (spore suspensions) were administered to each treatment group [29]. Prior to the administration of crude extracts and spore suspensions, the body weights of the mice were measured using analytical balance to prepare appropriate doses [29,38,39]. The animals in the experimental groups (n=5) were administered with four distinct concentrations of spores (1× 10^6^ ,1× 10^7^, 1× 10^8^ and 1× 10^9^ spores/mL) and four distinct concentrations of crude extracts (600, 1200, 2000 and 5000 mg/kg bw). A total of five female mice received each of the tested doses. Mice in the control group received the same volume of 0.9% NaCl [40].

### 2.8 Acute oral toxicity

Single-dose acute oral toxicity was evaluated based on OECD Guidelines [36,41]. The general behavior of mice and signs of toxicity (hypo-activity, breathing difficulty, tremors, and convulsion) were continuously observed for 1 h after oral treatment, after which these signs were intermittently observed for 4 h as well as over a period of 24 h [17,36]. Attention was given to signs of tremors, convulsions, salivation, diarrhea, lethargy, sleep and coma. During the subsequent post-dosing period (14 days), the animals were observed at least once per day. All of the observations were systematically recorded, with individual records maintained for each animal. Body weights were measured at the initiation of treatment, as well as 7 and 14 days after administration [37]. The LD_50_ value was determined according to the Dragstedt and Lang method described by El Allaoui, *et al*. [42].

### 2.9 Statistical analysis

Prism software (version 8.0; GraphPad, Inc., San Diego, CA) was used to perform the statistical analysis. Comparisons between the control and treatment groups were conducted using one-way analysis of variance (ANOVA), followed by a Tukey post-hoc test. All of the results were presented as the mean value ± standard error of the mean; the threshold for statistical significance was set at p≤0.05.

## 3. Results

### 3.1 Effect of temperature and pH on biocontrol agent mycelial growth

The investigated *Trichoderma* species showed optimum growth at 28°C on both minimal and yeast extract agar media, with neither species growing at 37°C. They were able to grow at all of the tested pH values (ranging from 2 to 9) at 28°C, with optimum growth observed at pH 4. It should be stated that both biocontrol agents were able to grow at pH 7, which is a precondition of growth within the human body.

### 3.2 Toxic effects of secondary metabolites against F. xylarioides

Methanolic crude extracts from both *Trichoderma* species amended with PDA medium were capable of inhibiting the mycelial growth of *F. xylarioides* (*p*≤0.05). *Trichoderma longibrachiatum* AU158 showed the highest defined level of *in vitro* antagonistic activity in both seeded agar assay and agar diffusion assay methods (Figures 1A, B and C). ANOVA analysis from seeded agar assay method revealed statistically significant (*p* ≤ 0.05) differences in the mycelial growth inhibition of *F. xylarioides* at different crude extract concentrations, with inhibition percentages ranging from 38 to 66.2% (*T. longibrachiatum* AU158) and 20 to 50.2% (*T. asperellum* AU131) (Figure 1C). Moreover, the crude extracts from *T. longibrachiatum* AU158 at maximum concentrations incited a change in the colony morphology of the pathogen with significant reduction in growth. The mean inhibitory effect against *F. xylarioides* restricted almost completely in plates as compared to the control, *F. xylarioides*, grown alone (Figure 1A). The agar diffusion test of the crude extracts from both *T. asperellum* AU131 and *T. longibrachiatum* AU158 were the most effective at a concentration of 200 μg/mL, causing 62.5%, and 74.3% inhibition, respectively (Figure 1B). The inhibition was not visible on the test pathogen at a concentration ≤50 mg/mL. The radial growth inhibition of *F. xylarioides* in both bioassay methods are attributed to inhibitory secondary metabolites released by bioagents through competition, mycoparasitism and production of cell wall degrading enzymes.

**Figure 1.**
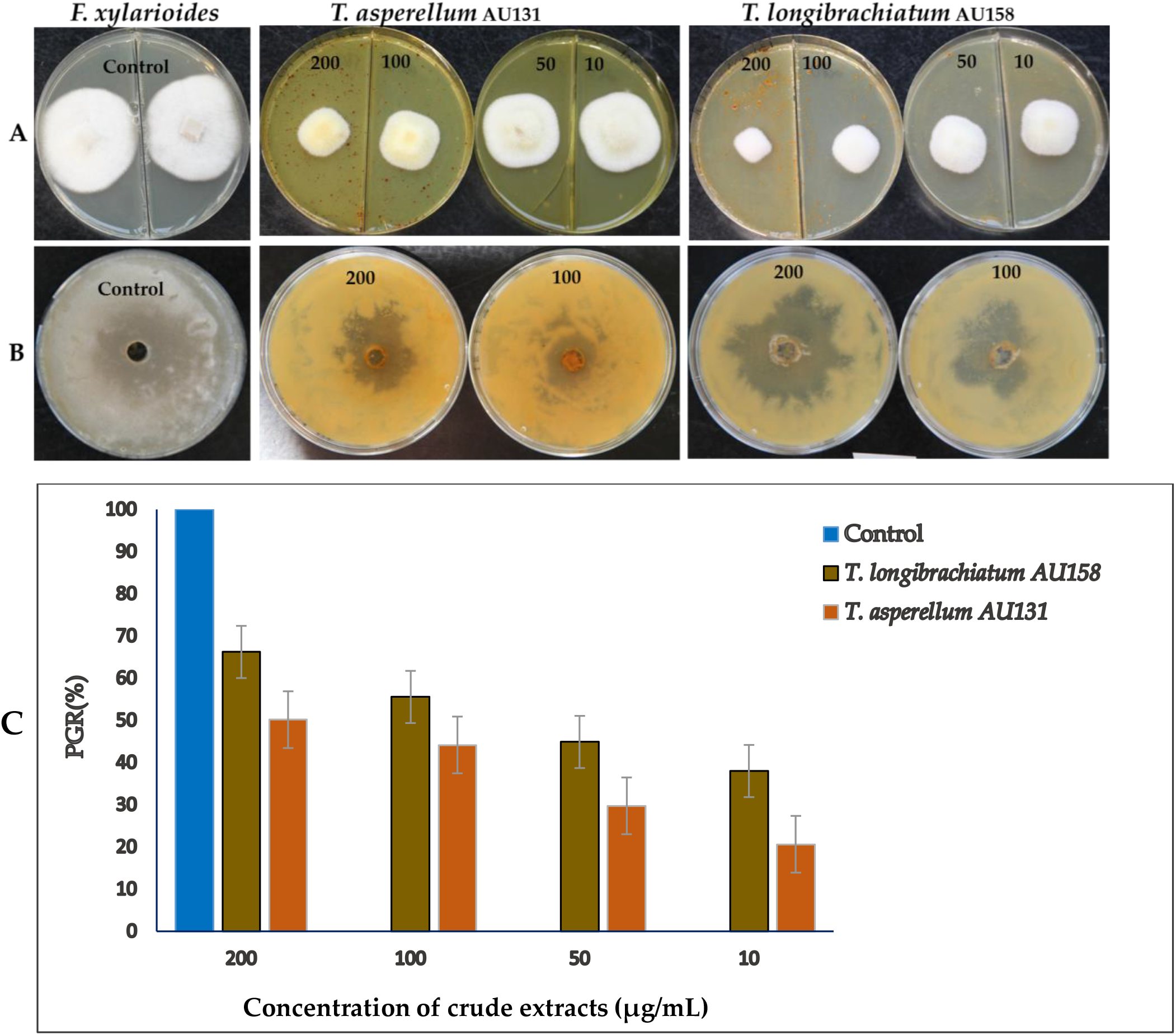
Bioassay activity of methanolic crude extracts of *T. asperellum* AU131 and *T. longibrachiatum* AU158 against *F. xylarioides*. **A** and **C**) Seeded agar assays: The picture shows the antifungal activity at different concentrations (200, 100, 50 and 10 μg/mL of crude extracts amended with PDA medium) and **B**) Agar diffusion assays (200 and 100 μg/mL of crude extracts).

### 3.3 GC-MS identification of secondary metabolites

A GC-MS analysis of the methanol crude extracts of both BCAs revealed the presence of 23 volatile secondary metabolites when the results were compared to the National Institute of Standard and Technology (NIST) library. The chromatogram illustrated that various volatile metabolites were present in the analyte, with these compounds then identified based on molecular weight, retention time, and molecular formula. Typical chromatograms and mass spectra of the compounds identified from the crude extracts of *T. asperellum* AU131 and *T. longibrachiatum* AU158 were shown in Figures 2A and 2B, respectively. The identified compounds were classified as alcohols, acids, sesquiterpenes, aromatics, ketones and esters according to the chemical class (Table 1). The compounds most commonly identified from the *T. asperellum* AU131 crude extract were, cis-13-Octadecenoic acid (21.48%), 2,4-bis(1,1-dimethylethyl)-Phenol (17.34%), *n* hexadecanoic acid (15.6%), hexadecanoic acid ethyl ester (14.14%), and linoleic acid ethyl ester (11.46%), while the compounds most commonly detected from the *T. longibrachiatum* AU158 crude extract were cis-13-Octadecenoic acid (23.48%), 2,4-bis(1,1-dimethylethyl)-Phenol (18.5%), ethyl oleate (17.83%) and *n*-hexadecanoic acid (16.72) (Table 1).

**Figure 2.**
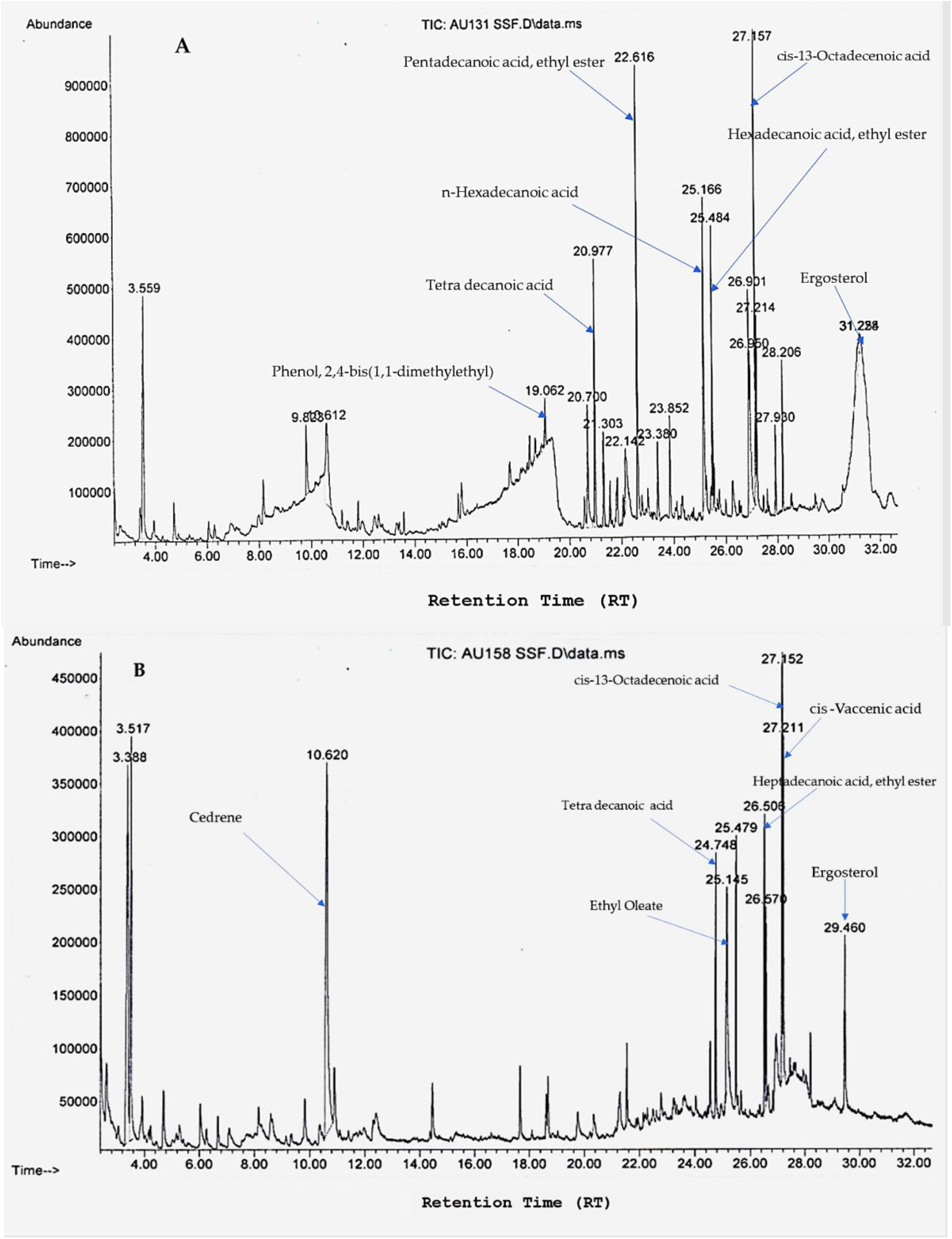
Chromatograms of the volatile organic compounds identified from *T. asperellum* AU131 (A) and *T. longibrachiatum* AU158 (B) crude extracts.

**Table 1.**
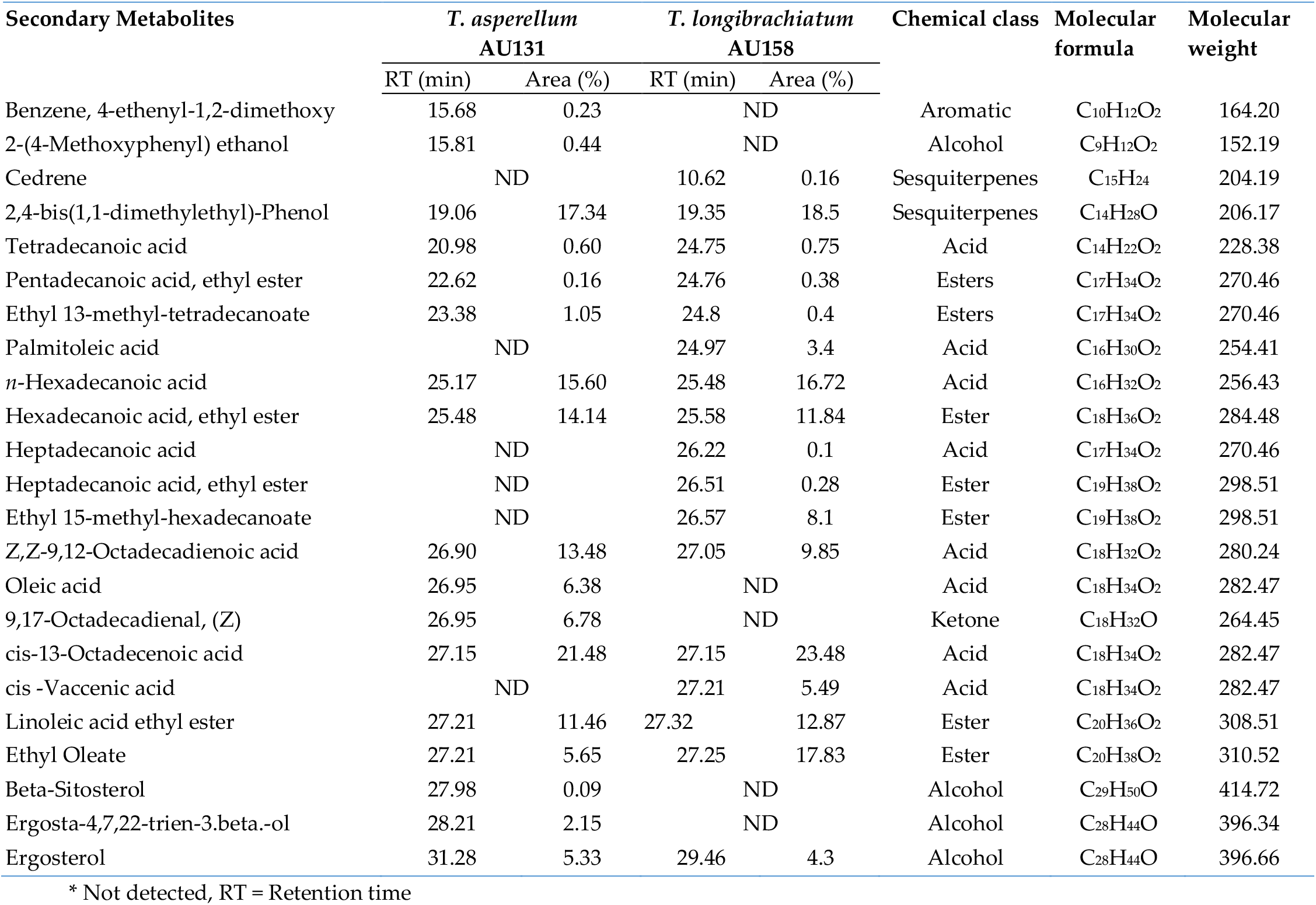
Volatile organic compounds of *T. asperellum* AU131 and *T. longibrachiatum* AU158 and identified by GC-MS.

### 3.4 Acute toxicological evaluation

The survival rate in the assay was 100%. In the acute toxicity study, crude extract doses under 5000 mg/kg bw and spore doses under 1 × 10^9^ spores/mL did not cause any mortality or signs of toxicity in female mice over the observed period. No obvious clinical signs, including hair loss, scabbing, soft or mucoid feces, and lowered defecation, were noted in the study animals. All of the experimental animals showed normal behavior 14 days after administration, which demonstrates that the metabolites produced by the studied species as well as direct administration of live cells are safe for mice. Furthermore, the administration of metabolites or live cells did not cause any inflammatory or enterotoxigenic effects that could have induced gastroenteritis, e.g., diarrhea. On the other hand, it was not possible to calculate the mean lethal dose (LD_50_) because the administered doses did not cause death in any of the mice. In other words, the LD_50_ test results concerning methanolic extracts and spore suspensions mean that the metabolites and *Trichoderma* species do not cause any lethal effects in mice at doses below 5000 mg/kg bw and 1 × 10^9^ spores/mL, respectively.

### 3.5 Effect of crude extracts and spore suspensions on body weight of mice

Animals in the treatment group, i.e., which were administered with crude extracts or live cells, did not show any apparent changes in body weight, with the values similar to what was observed in the control group, which were administered with a 0.9% NaCl solution. The results illustrated in Figures 3A and 2B demonstrate that mice in both the treatment and control groups showed a gradual increase in body weight over the study period. The results presented in this study showed that the oral administration of either crude extracts or cellular suspensions did not cause any significant (p>0.05) changes in the weight of mice. Furthermore, the body weights of animals in the treatment group did not noticeably differ from the body weights of animals in the control group. This suggests that the crude extracts and spores of the two tested *Trichoderma* species neither support weight loss nor stimulate weight gain.

**Figure 3.**
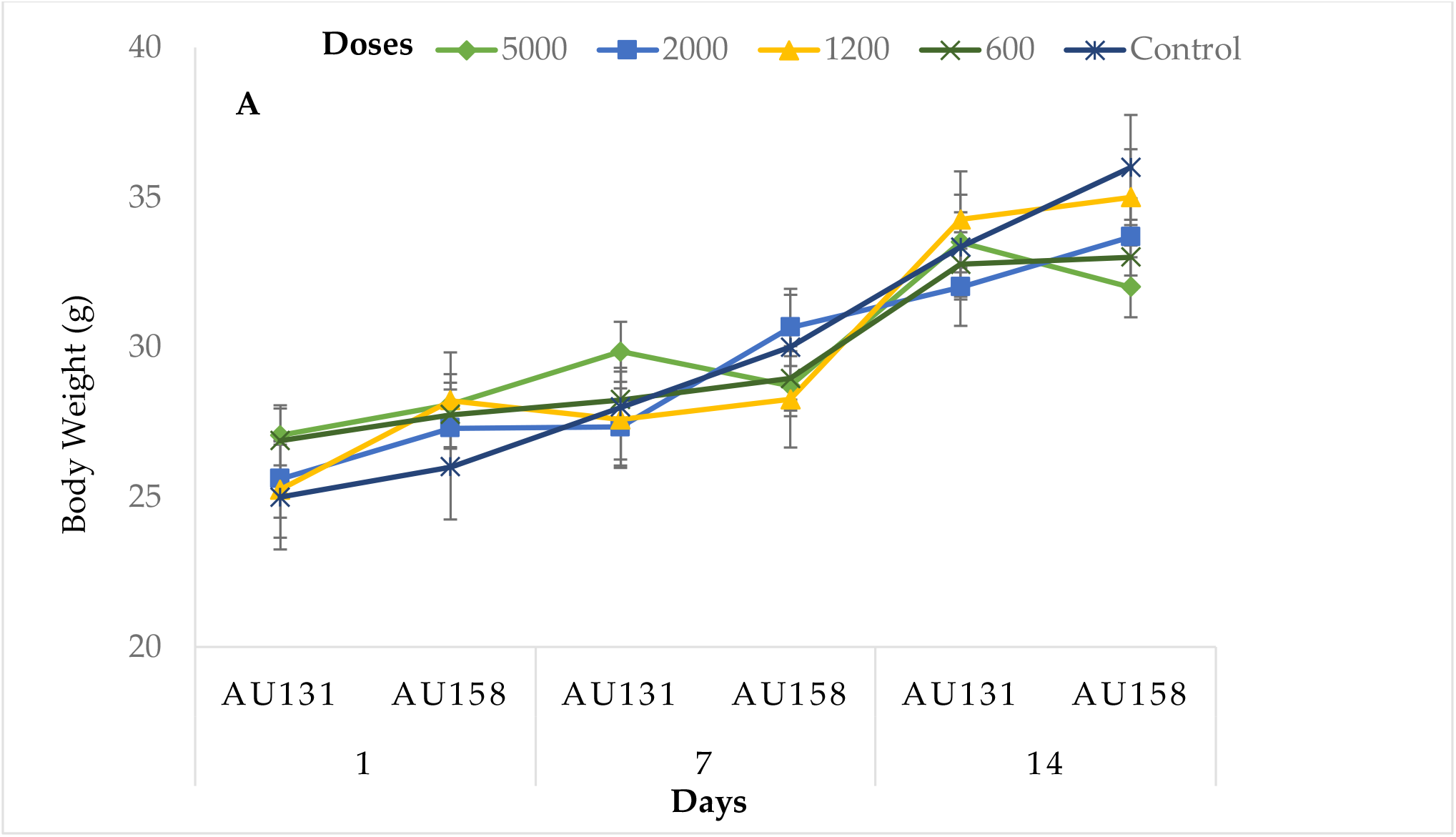
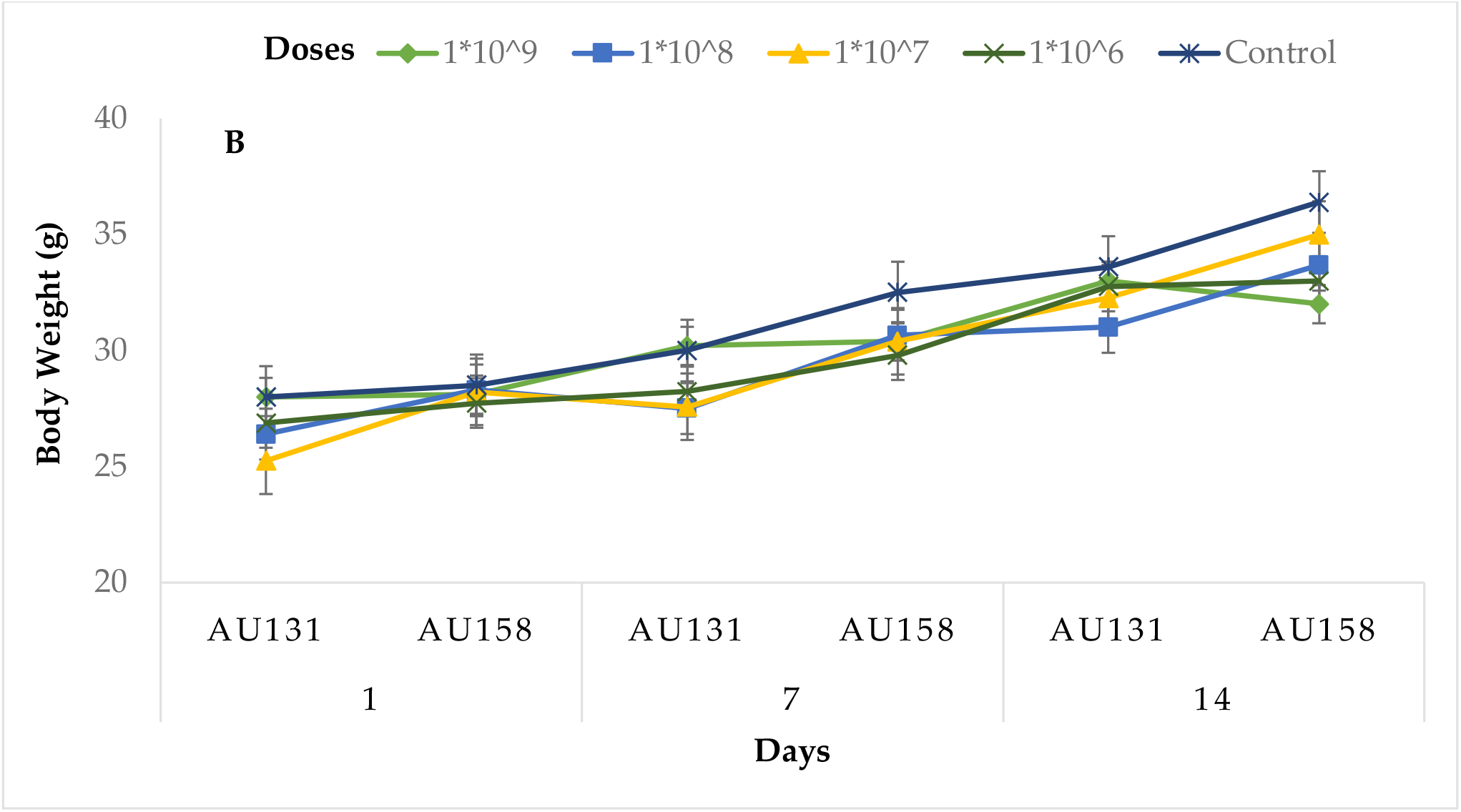
Changes in the body weights of mice administered with crude extracts and spore suspensions from *Trichoderma* species. (A) Mice that were administered with methanolic crude extracts and (B) mice that were administered with spore suspensions. Each point represents the mean value ± SD (n = 5).

## 4. Discussion

In the previous study, we formulated *T. asperellum* AU131 and *T. longibrachiatum* AU158 based biofungicides under SSF to control *F. xylarioides*, the causative agent of CWD in Ethiopia [16]. The present study investigated the in vitro bioassay and acute toxicity of these biocontrol agents through short-term assays in animal models to ensure their safety in coffee farm applications. Most of the species involved in *Trichoderma* infections are considered as opportunistic pathogens [43], with the virulence factors including mycelial growth at 37°C and neutral pH, hemolytic activity and toxicity to mammalian cells [13,14], along with resistance to antifungal compounds. The results of the present study showed that neither *T. asperellum* AU131 nor *T. longibrachiatum* AU158 grew on minimal media at 37°C. However, both species were able to grow at physiological pH, which agrees with the reports of Antal, Kredics, Pakarinen, Dóczi, Andersson, Salkinoja-Salonen, Manczinger, Szekeres, Hatvani and Vágvölgyi [13] and Kredics, Antal, Szekeres, Manczinger, Dóczi, Kevei and Nagy [14]. More specifically, Antal, Kredics, Pakarinen, Dóczi, Andersson, Salkinoja-Salonen, Manczinger, Szekeres, Hatvani and Vágvölgyi [13] reported that *T. longibrachiatum* strains showed optimum growth at 30°C on both minimal and yeast extract agar media, while all of the strains were also able to grow at 40°C. Mikkola, *et al*. [44] reported that toxic *T. longibrachiatum* strains grew on malt extract agar (MEA) at 22°C and at 37°C. Hence, the concerns and risks associated with biopesticides can differ from one species to another, most likely requiring a case-by-case approach in risk assessment.

*Trichoderma* species are known to produce a wide range of bioactive secondary metabolites that are known to have antifungal, antibacterial, and toxic properties to control a wide range of phytopathogens, such as *Fusarium* species, *Botrytis cinerea, Pythium* species, *Rhizoctonia solani, Sclerotinia sclerotiorum*, and *Ustilago maydis* [8,21,33]. In the present study, the results of the antifungal activity of crude extracts obtained from both *T. asperellum* AU131 and *T. longibrachiatum* AU158 showed that the extracted secondary metabolites inhibited the growth of *F. xylarioides* in the seeded agar and agar diffusion assays, as shown in Figures 1A and 1B. Both extracts were shown to have inhibitory activity, however, the extract of *T. longibrachiatum* AU158 was more effective, as it was able to inhibit at the minimum concentration (10 mg/mL) in seeded agar assay. Many species of *Trichoderma*, such as *T. harzianum, T. viride, T. hamatum, T. atroviride*, and *T. virens*, have been reported to be effective in control of a wide range of soil borne plant pathogens [45,46]. However, the biocontrol efficacy varies with *Trichoderma* species and target plant diseases [47]. A significant advancement from the present study is the finding that the SSF and crude extract of the AU131 and AU158 strains provided a significant inhibitory effect on *F. xylarioides*, a causative agent of coffee wilt disease. Our findings suggest that both strains can be considered as a biocontrol agent in the effort of using alternative approaches to control coffee wilt disease.

Analysis of the GC-MS chromatograms of the crude extracts from both *T. asperellum* AU131 and *T. longibrachiatum* AU158 revealed the production potential secondary metabolites capable of inhibiting the growth of *F. xylarioides*. The analysis showed that both BCAs produced large quantities of volatile organic compounds (VOCs) that were identified as alcohols, acids, esters, sesquiterpenes and ketones. This wide range of VOCs emitted by both *Trichoderma* species is in line with the diverse VOCs produced by *Trichoderma* species reported in previous studies [33,48]. The volatile secondary metabolites identified from crude extracts of the two investigated *Trichoderma* species (*T. asperellum* AU131 and *T. longibrachiatum* AU158) also agreed with what has previously been presented in the literature [33-35]. The fungal production of VOCs is a dynamic process in that it is directly affected by both genetic and environmental factors, which include community composition, substrate, temperature, moisture level, and pH [49-52]. The identified metabolites play important roles in mycoparasitic interactions as well as induced systemic resistance (ISR) in plants via the upregulation of jasmonic acid and salicylic acid synthesis [53]. For instance, 2,4-bis(1,1-dimethylethyl)-phenol (2,4-DTBP) was found to be effective against the agriculturally important root-rot fungus; *Fusarium oxysporum* based on the inhibition of spore germination and hyphal growth [54]. During fungal spore germination, 2,4-DTBP completely inhibited germination by preventing the emergence of a normal germ tube, eventually leading to abnormal branching and swelling of the hyphae [54].

As GC-MS is the first step towards profiling the metabolites present in crude extracts, the present study showed presence of five major VOCs from both BCAs viz. cis-13-Octadecenoic acid, 2,4-bis(1,1 dimethylethyl)-Phenol, *n*-hexadecanoic acid, ethyl oleate and hexadecanoic acid ethyl ester (Table 1). Most of the metabolites identified in this study are widely used as antimicrobials, food additives, cancer drugs, herbicides and pesticides. Specifically, cis-13-Octadecenoic acid and *n*-hexadecanoic acid are reported to have anti-inflammatory, cancer preventive and hepatoprotective properties [55]. The other identified linoleic acid ester and Z,Z-9-12-Octadecenoic acid also have anti-inflammatory, antiandrogenic, and anemiagenic properties [56]. Moreover, two of the identified volatile compounds – tetradecanoic acid and *n*-hexadecanoic acid methyl ester – are antioxidants [57]. Since there are no complete toxicity reports for all the secondary metabolites from *Trichoderma* species, we tested their toxicological levels using animal models (mice). We used methanolic crude extracts to assess the toxicological risks associated with a particular BCA rather than using purified commercial metabolites as per the guidelines of the OECD.

Crude extracts and spore suspensions were first administered by oral gavage, since this is the most common route of human exposure. Thus, the acute toxicity assays were performed to monitor the harmful effects of an biocontrol agents to the organism following single or short-term exposure [58]. The performed experiments mainly evaluated mortality, changes in behavior, body weight, and other characteristics relative to the overall well-being of mice. In the present study, the *in vivo* toxicity evaluation of methanolic crude extracts and spore suspensions of *Trichoderma* species did not reveal any mortality among the Swiss albino mice; this suggests that the extracts and spore suspensions are not deadly to mammals. A report by the European Food Safety Authority EFSA [59] indicated that *T. harzianum* Rifai strain T-22 and *T. asperellum* T25 did not result in adverse effects following the oral, intratracheal, subcutaneous and intravenous administration of doses ranging from 6.4 × 10^6^ to 1.5 × 10^7^ colony forming unit (cfu)/animal. Moreover, the report indicated complete clearance of the spore suspension within two days of oral administration, a dramatic decrease in levels in the lungs by day 21 post-intratracheal administration, and a marked decrease in the number of cfu from the colonized organs following intravenous administration. No signs of pathogenicity were observed for any of the routes of exposure, which is in agreement with the results of the present study.

However, crude extracts of various *T. longibrachiatum* isolates have been reported to contain heat-resistant substances that inhibit the motility of boar spermatozoa and quench the mitochondrial transmembrane potential (ΔΨm) of sperm cells at low concentrations [60]. Several clinical isolates originally identified based on their morphological characters were recently reidentified by sequence-based molecular techniques as *T. longibrachiatum*, which provided the most frequently occurring clinical etiological agent within the genus *Trichoderma* [61,62]. Thus, the biotechnological and agricultural application of *T. longibrachiatum* strains should be carefully monitored to minimize possible health risks.

Crude extracts from *T. asperellum* AU131 showed no toxicity up to concentrations of 5000 mg/kg bw. Moreover, acute oral toxicity studies conducted with both *T. asperellum* AU131 and *T. longibrachiatum* AU158 cellular suspensions did not result in treatment-related adverse effects across doses ranging from 1 × 10^6^ to 1 × 10^9^ spores/mL. This was supported by previous observations that the oral administration of crude extracts and cellular suspensions representing various fungal BCAs demonstrated very low toxicity in rodents and ruminants [26,63]. Moreover, a report from the USA Environmental Protection Agency [25] demonstrated that *Trichoderma asperellum* strain ICC 012 is non-toxic to rats at a dose of 2000 mg/kg bw and 4.2 × 10^9^ cfu/g. The product formulated from this species contains 99.9% *w/w Trichoderma asperellum* strain ICC 012 as the active ingredient, with a minimum and nominal content of 1 × 10^9^ and 2.5 × 10^9^ cfu/g dry weight), respectively. The report clearly stated that exposure to this species does not cause any acute, sub-chronic, chronic, immune, endocrine, or non-dietary issues.

On the other hand, the oral median lethal dose (LD_50_) could not be calculated since the administered doses did not cause any death to the mice. These results suggest that the lethal doses for both *T. asperellum* AU131 and *T. longibrachiatum* AU158 crude extracts exceed 5000 mg/kg bw, which represents the highest reference dose [17]. In this regard, crude extracts may be considered safe at the tested levels, i.e., 5000 mg/kg bw and below. Changes in body weight is an important index for evaluating the toxicity of bioformulated products [64]. In the present study, both the treatment and control groups demonstrated a gradual increase in mean body weight. However, the difference in weight gain between the control and treatment groups was statistically insignificant (p≥0.05).

Moreover, the study evaluate the acute toxicity of *T. asperellum* AU131 and *T. longibrachiatum* AU158 using Swiss albino mice and GC-MS to identify volatile metabolites of methanolic extracts. However, non-volatile metabolites produced by *Trichoderma* species and showing antimicrobial activity such as gliotoxin, peptaibols, gliovirin, polyketides, and pyrones were not addressed in this study since they are not detected by GC-MS. Thus, further study is under progress to quantify and assess their toxicity related risks by using untargeted Liquid chromatography–high resolution mass spectrometry (LC–HRMS).

## 5. Conclusion

The two *Trichoderma* species tested in this study, *T. asperellum* AU13 and *T. longibrachiatum* AU158, inhibited the growth of *F. xylarioides* in seeded agar assay and agar diffusion assay methods. In particular, the extract of *T. longibrachiatum* showed the highest inhibition activity and was active even at a low concentration. These results highlight the potential use of these two *Trichoderma* species as antagonists in biocontrol of coffee wilt disease. On the other hand, based on the results of this study, it can be concluded that the tested biocontrol agents (*T. asperellum* AU131 and *T. longibrachiatum* AU158), along with crude extracts and spore suspensions, were neither toxic nor pathogenic to Swiss albino mice across the tested range of concentrations, respectively. Both *Trichoderma* species showed optimum growth at 28°C on minimal and yeast extract agar media, with neither species growing at 37°C. A GC-MS analysis of the methanol crude extracts from both bioagents revealed the presence of 23 VOCs that were classified as alcohols, acids, sesquiterpenes, ketones and aromatic compounds. However, further study is required to verify the toxicity level of each metabolites.

## Author Contribution

Conceptualization, A.M and T.A; methodology, A.M., T.A and N.M.; formal analysis, A.M.; investigation, A.M., and T.T.; resources, T.A. and R.R.V.; data curation, T.T., T.A., N.M. and R.R.V.; writing—original draft preparation, A.M.; writing—review and editing, T.A., T.T., N.M. and R.R.V.; supervision, T.A., N.M. and R.R.V.; project administration, T.A; funding acquisition, T.A., and R.R.V. All authors have read and agreed to the published version of the manuscript.

## Funding

This research project was supported by the Ethiopian Biotechnology Institute (EBTi), Ethiopia, under the program of Biotechnology Product Development Research Grant.

## Institutional Review Board Statement

The name of the ethics committee is the Laboratory Animal Ethics Committee, College of Natural and Computational Sciences Institutional Review Board (CNCS-IRB), approved the implementation of the experiment with the ethical approval code IRB/44/2020, and the date of approval was 07 January 2020.

## Acknowledgments

We would like to thank EBTi for funding this research project under the theme “Bio-fungicide production for CWD control in Ethiopia”. We also thank Department of Microbial, Cellular and Molecular Biology and Department of Chemistry of Addis Ababa University (AAU), Ethiopia, for the laboratory facilities. Special thanks goes to the Department of Plant Breeding of the Swedish University of Agricultural Sciences (SLU), Sweden, for valuable support to the project. RV was supported by the Swedish Research Council for Environment, Agricultural Sciences and Spatial Planning (FORMAS) (grant number 2019-01316) and the Swedish Research Council (grant number 2019-04270).

## Informed Consent Statement

Not applicable.

## Conflicts of Interest

The authors declare no conflict of interest.

## Notes

### Competing Interest Statement

The authors have declared no competing interest.

